# Loss and recovery of transcriptional plasticity after long-term adaptation to global change conditions in a marine copepod

**DOI:** 10.1101/2020.01.29.925396

**Authors:** Reid S. Brennan, James A. deMayo, Hans G. Dam, Michael Finiguerra, Hannes Baumann, Melissa H. Pespeni

**Author notes:** Authors contributed equally to this work (Reid S. Brennan, James A. deMayo).

## Abstract

Adaptive evolution from standing genetic variation and physiological plasticity will fuel resilience in the geologically unprecedented warming and acidification of the earth’s oceans. For marine animals, however, we have much to learn about the mechanisms, interactions, and costs of adaptation. Here, using 20 generations of experimental evolution followed by three generations of reciprocal transplantation, we investigate the relationship between adaptation and plasticity in the marine copepod, *Acartia tonsa*, in future greenhouse conditions (high temperature, high CO_2_). We find highly parallel genetic adaptation to greenhouse conditions in genes related to stress response, gene expression regulation, actin regulation, developmental processes, and energy production. However, reciprocal transplantation showed that genetic adaptation resulted in a loss of transcriptional plasticity, reduced fecundity, and reduced population growth when greenhouse animals were returned to ambient conditions or reared in low food conditions, suggestive of genetic assimilation after 20 generations of adaptation. Despite the loss of plasticity at F21, after three successive transplant generations, greenhouse-adapted animals were able to match the ambient-adaptive transcriptional profile. Concurrent changes in allele frequencies and erosion of nucleotide diversity suggest that this recovery occurred via adaptation back to ancestral conditions. These results demonstrate the power of experimental evolution from natural populations to reveal the mechanisms, timescales of responses, consequences, and reversibility of complex, physiological adaptation. While plasticity facilitated initial survival in global change conditions, it eroded after 20 generations as populations genetically adapted, limiting resilience to new stressors and previously benign environments.

## Main Text

Though global conditions are changing at a geologically unprecedented rate and the frequency of extreme events is increasing due to human activities ^1^, species can use genetic adaptation, physiological plasticity, or both to persist ^2–4^. Genetic adaptation, heritable genetic change that improves the mean fitness of a population in an environment ^5^, enables resilience by shifting the mean phenotype of the population to tolerate different conditions. Selection can shift population genetic variation over short time scales, as few as one to four generations, changing population phenotypes to improve mean fitness in response to an extreme change in the environment ^6–9^. Conversely, plasticity allows a single genotype to generate different phenotypes in response to the environment ^10^. Considering rapid changes in environmental conditions, adaptive plasticity enables organisms to maintain fitness across environments ^11^ through underlying phenotypic changes, such as shifts in metabolism or gene regulation ^12^. However, the interaction and relative contributions of adaptation and plasticity to population persistence in rapid environmental change still remains a critical area of investigation ^13–16^.

Theoretical models predict that plasticity may increase when a population is exposed to a novel environment, but should decline across longer timeframes ^17^. Indeed, a reduction of plasticity appears to be common in natural and experimental empirical systems ^18–22^. However, this experimental work has also revealed faster losses of plasticity during rapid adaptation than theory would predict. For example, Waddington showed that a high temperature induced plastic phenotype, crossveins in *Drosophila* wings, can become fixed, or assimilated, with 20 generations of artificial selection, reducing plasticity and the environmental responsiveness of a trait ^23^. Under climate change conditions, populations will experience strong selection to persist in stressful environments. Thus, it is essential to understand how selection in these novel conditions may alter or reduce the plasticity of natural populations.

Experimental evolution is a particularly powerful approach to understand resilience as organisms can be exposed to specific selective pressures over multiple generations ^24^. From this, one can identify both the adaptive potential and the mechanisms of adaptation of a population, directly providing insight into how a population may cope with changing climates ^25^. Experimental evolution in fruit flies ^26, 27^, copepods ^20^, marine algae ^28^, nematodes ^21^, mice ^29^, and others ^30^ have identified phenotypic and/or genomic responses to selection and have demonstrated that strong selection can rapidly shift or drive a loss of plasticity. The majority of these studies have leveraged model systems and found that when selection regimes are static, plasticity tends to be lost (but see ^27, 31^). Thus, this raises the concern that the strong selection driving rapid adaptation to climate change conditions may also purge phenotypic plasticity from populations, for example, as seen in marine to freshwater transitions in copepods in the wild ^18^. Unfortunately, in non-model metazoans, experimental evolution experiments tend to be restricted to a small number of generations with a single selective pressure and few have measured the ability of organisms to match their phenotype to a multiple stressor environment across multiple generations of experimental evolution and multiple generations of reciprocal transplant. Such experiments could test the effects of long-term adaptation and the potential for recovery of plasticity. Incorporating this multigenerational perspective is essential to better understand if organisms can shift phenotypes following rapid environmental change.

The use of outbred natural populations in evolution experiments is a powerful approach for revealing the potential contributions of standing genetic variation and plasticity, particularly in context of future global change conditions ^32, 33^. Species that live in dynamic environments, that have distributions that span a wide range of environmental conditions, and that have high dispersal capacity are predicted to have the greatest capacity to respond to rapid environmental change due to their high levels of standing genetic variation and physiological plasticity ^3, 34^. Many marine species in particular live in environments that vary across time and space and have life histories that promote dispersal ^35–37^. This is true of copepods, the most abundant marine metazoan ^38^, which have plasticity and genetic variation for adaptation to global change conditions ^20, 39–42^. However, there is currently limited empirical evidence of the capacity for and genetic bases of adaptation to global change conditions over longer timescales (e.g., greater than 2 generations) in natural populations of copepods, or any marine metazoan ^32, 33^ (but see ^43^), and the relationship between plasticity and genetic adaptation is only beginning to be understood in marine systems ^31, 44, 45^. Given their ecological importance and high levels of genetic variation and plasticity, copepods are an ideal model to begin to disentangle the interplay of genetic and plastic responses of natural populations of metazoans to global change conditions.

We experimentally evolved the globally distributed, foundational copepod species, *Acartia tonsa,* to ambient and global change conditions to determine the effects of long-term adaptation on plasticity. As a dominant prey item for forage fish, *A. tonsa* serves as a critical link in the marine food web, supporting economically important fisheries ^46^ and as major phytoplankton grazers contribute to the storage of atmospheric CO_2_ and thus mediate marine biogeochemical cycles ^47^. Replicate cultures (4 replicates per condition, ∼4,000 individuals per replicate) of *A. tonsa* were subjected to 20 generations of selection in ambient (AM: 400 ppm *p*CO_2_ and 18°C) and combined high CO_2_ × temperature (greenhouse, GH: 2000 ppm *p*CO_2_ and 22°C) conditions followed by three generations of reciprocal transplantation of both lines to opposite conditions (Fig. 1). These conditions were chosen as they represent present-day and a worst case, yet realistic scenario ^48, 49^; in the coming century, global mean ocean surface temperatures will increase by 2-4 °C ^50^ and oceanic CO_2_ concentrations will potentially reach 2000 ppm by the year 2300 ^51^. We denote evolved lines with their abbreviation and the environmental condition during transplant as a subscript (i.e., AM_GH_ = ambient line transplanted to greenhouse conditions). RNA was collected from pools of 20 adults from each of four replicate cultures at the first, second, and third generation after transplant for both experimentally evolved and transplanted lines (Fig. 1). We estimated allele frequencies at 322,595 variant sites with at least 50× coverage across all samples (mean coverage 174×) and quantified transcript abundance for an average 24,112 genes.

**Figure 1:**
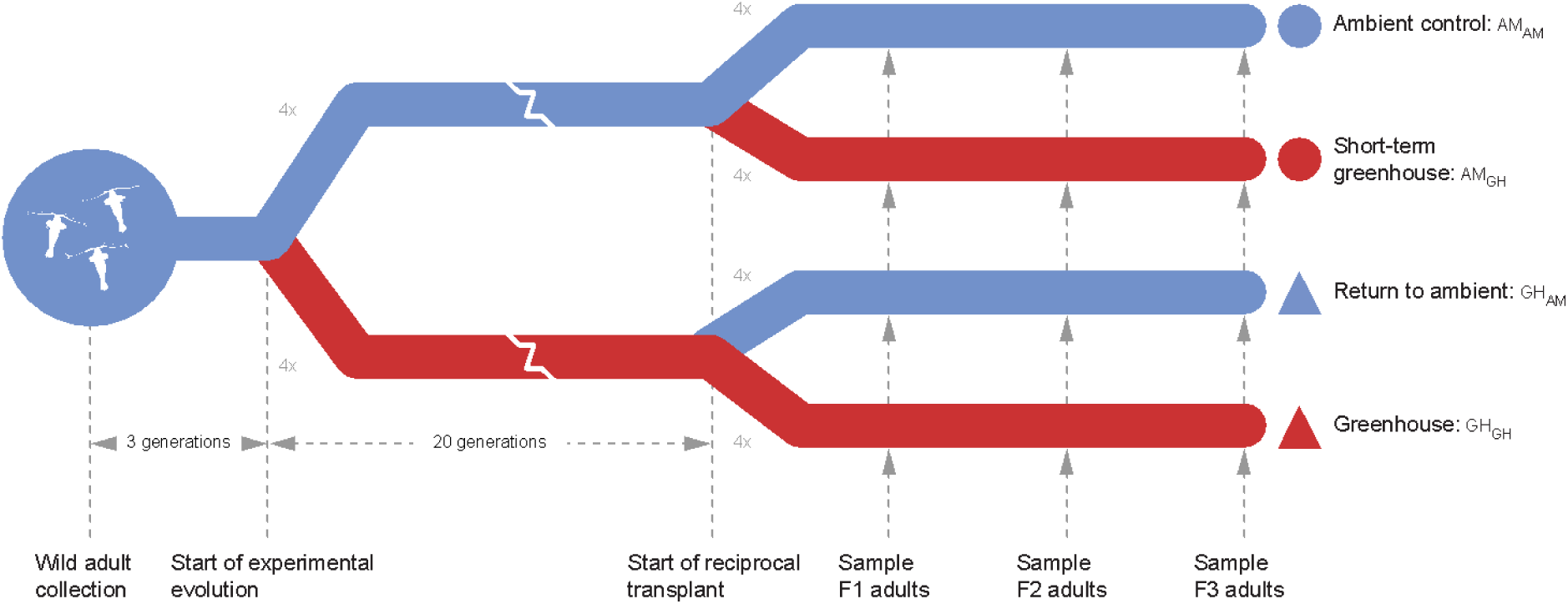
Schematic of the experimental design. Blue lines are ambient (AM) pH and temperature that represent current conditions (*p*CO_2_: 400 ppm; temperature: 18°C). Red lines are simulated future greenhouse (GH) conditions (*p*CO_2_: 2000 ppm; 22°C). Adult *Acartia tonsa* were collected from the wild and reared in the lab for three generations. Six hundred laboratory-acclimated adults seeded each of four replicates at AM and GH conditions where they were reared for 20 non-overlapping generations. At generation 20, each replicate was split in two and transplanted into the same conditions as the previous 20 generations (AM_AM_, GH_GH_) and to the opposite condition (AM_GH_, GH_AM_). These transplanted lines were reared for 3 additional generations and sampled for life history traits at the first generation and genomics at the end of each generation.

We address three major questions in this study. 1) Are copepods able to adapt to global change conditions and, if so, what are the allelic and gene expression mechanisms underlying adaptation? 2) To what extent is genetic adaptation versus transcriptional plasticity required to respond to environmental change? Specifically, does adaptation result in a loss of plasticity in a marine metazoan? 3) Does adaptation result in trade-offs among life-history traits revealed through reciprocal transplant and in low food stress conditions? We predicted that this coastal copepod would have genetic and plastic mechanisms to respond to greenhouse conditions and that long-term adaptation would reduce physiological plasticity. Our results support these predictions, revealing the molecular underpinnings of the adaptive mechanisms, and demonstrate a loss and recovery of transcriptional plasticity at the expense of adaptive genetic variation.

## Results and discussion

Genetic and transcriptional variation consistently diverged after 20 generations in greenhouse versus ambient conditions (Fig. 2, S1, S2), suggesting directional selection in response to global change conditions. A total of 17,720 loci (5.5% of 322,595 loci with >50X coverage) showed consistent allele frequency divergence (GH_GH_ versus AM_AM_; Cochran–Mantel–Haenszel test FDR corrected significance threshold: 5.17e-8). Given the likely polygenic nature of adaptation to greenhouse conditions and that polygenic traits tend to be genetically redundant ^52, 53^, the identified adaptive genetic variation would be prone to false negatives due to the requirement that adaptation be parallel in all replicates. Conversely, linkage disequilibrium contributes some number of false positives when neutral loci are located near the selected loci ^54^. Of potential candidate genes driving adaptation (Fig. S2), one gene, NADH-ubiquinone oxidoreductase 49 kDa subunit (NDUFS2), was an extreme outlier for divergence in allele frequencies, 77% change, which is 72% greater than the global average (5% change in allele frequency; Fig. S3). NDUFS2 is a nuclear-encoded, core subunit of Complex 1 of the mitochondrial electron transport chain (ETC) that has been shown to be inhibited by heat stress ^55, 56^. In addition, mitochondrial function has been linked to variation in thermal tolerance among populations of the intertidal copepod *Tigriopus californicus* ^57^. The three substitutions identified here were all non-synonymous, including a non-polar Isoleucine and polar Asparagine, in linkage disequilibrium (Fig. S2), and in a single alpha helix of the ETC subunit. This alternative NDUFS2 haplotype targeted by selection could improve energy production under heat stress for global change-adapted copepods, a hypothesis worth future functional investigation ^58^.

**Figure 2.**
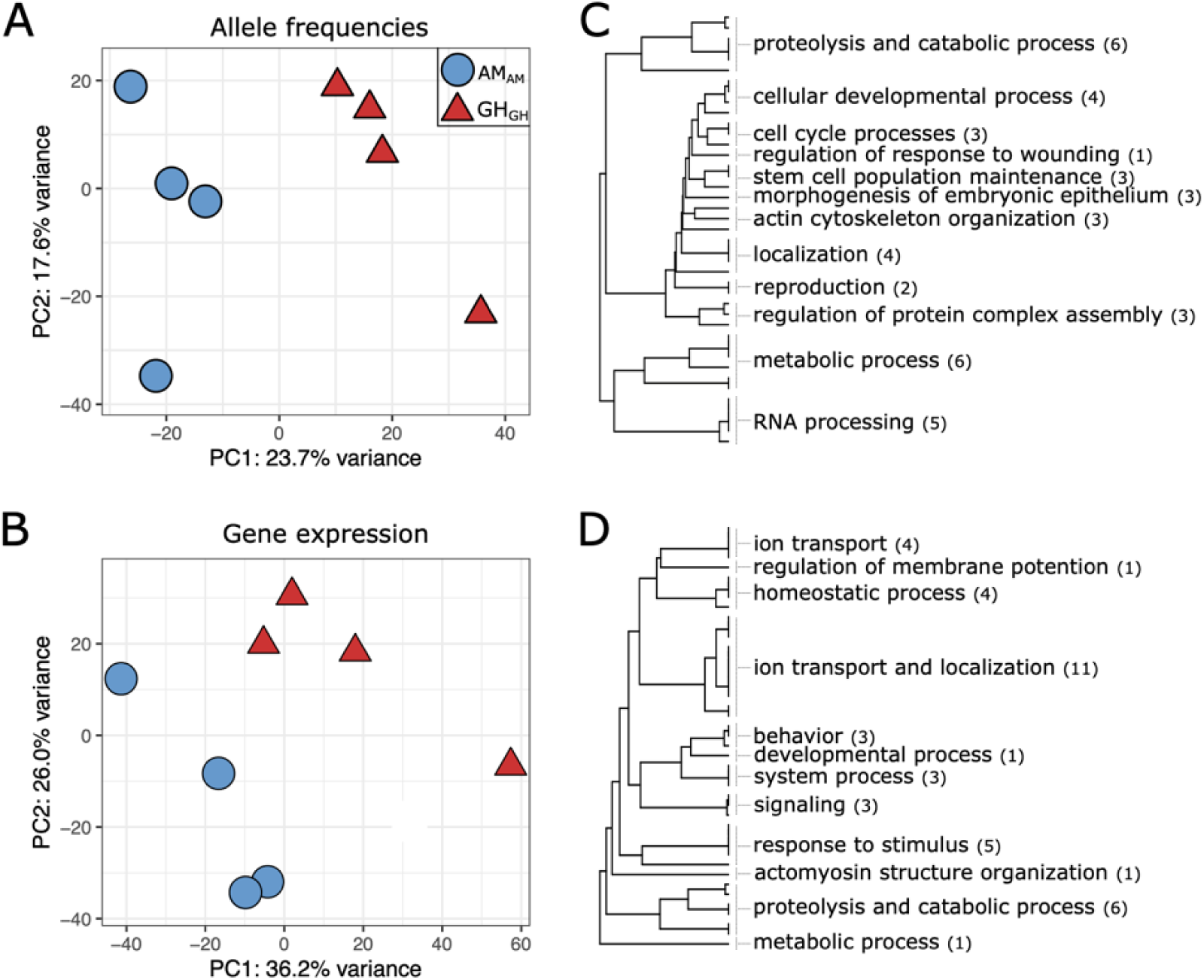
Allele frequency and gene expression divergence after 20 generations of selection. Principal component analysis of (A) genome-wide variation in allele frequencies (322,595 SNPs) and (B) gene expression (24,927 genes) at the F1 generation. (C, D) Gene ontology enrichment results from Mann-Whitney U test using p-values from Cochran–Mantel–Haenszel tests (allele frequencies, C) and DESeq2 Wald tests (gene expression, D). Gene categories are collapsed for visualization purposes with the number of categories indicated in parentheses. See tables S1-2 and figs. S7-8 for full results.

A total of 1,876 genes (7% of the 24,927 genes surveyed) were differentially expressed between control and experimentally evolved replicate lines in their home environment, differences that could be driven by plastic and evolved mechanisms (GH_GH_ versus AM_AM_; *P* < 0.05). There was substantial overlap in the functional classes of genes that had evolved allele frequency and gene expression differences. Shared categories included response and detection of stress and stimuli, developmental processes, actin organization and regulation, and proteolytic processes (Fig. 2C, D; Tables S1, S2). These functions may underlie increased survival in warmer and more acidic conditions. For example, proteolysis is integral to the stress response ^59^ while actin modifications have been identified as targets of pH adaptation and resilience in other marine organisms ^9, 60^ and are linked to cytoskeleton maintenance that is responsive to low pH stress in copepods ^61^. Despite shared functions, there was little overlap in the specific genes that evolved allele frequency and gene expression differences (16%; Fisher’s exact test; *P* > 0.05), suggesting that adaptive divergence in protein function and gene expression targeted different, but functionally related genes ^62^. Similarly, there was no correlation between the degree of expression and allelic responses (Fig. S4; R^2^ = 0.002), suggesting no impact of potential allele-biased expression on estimates of allele frequencies from pools of sequenced RNA. However, we found strong allele frequency divergence in regulators of expression, predominantly in genes relating to RNA processing (Fig. 2C) and also ribosomal S6 kinase, a gene involved in the regulation of numerous transcription factors and translation (Fig. S2) ^63^. Taken together, these results suggest that regulators of gene expression were under selection and may have been responsible for the adaptive changes in gene expression under global change conditions, in accord with previous work in copepods ^41^.

We next quantified gene expression plasticity to test the prediction that rapid adaptation to a stressful environment would result in a loss of physiological plasticity in GH animals. We identified plastic changes in gene expression for GH and AM lines by comparing expression in the home environment after 20 generations to gene expression responses at the end of one generation in transplant conditions (AM_AM_ vs. AM_GH_ and GH_AM_ vs. GH_GH_; Fig. 3A). At the first generation of transplant, AM lines showed plastic expression responses for 4,719 genes (Padj< 0.05; 20% of genes), a 12.7 fold greater response than for GH lines (372 genes; 1.6% of genes). However, at the end of the second and third generation in transplant, this difference diminished due to the relative increase of GH gene expression changes (F2: 4.5%; F3: 4.1%) and decrease of AM changes (F2: 1.7%; F3: 6.3%).

**Figure 3.**
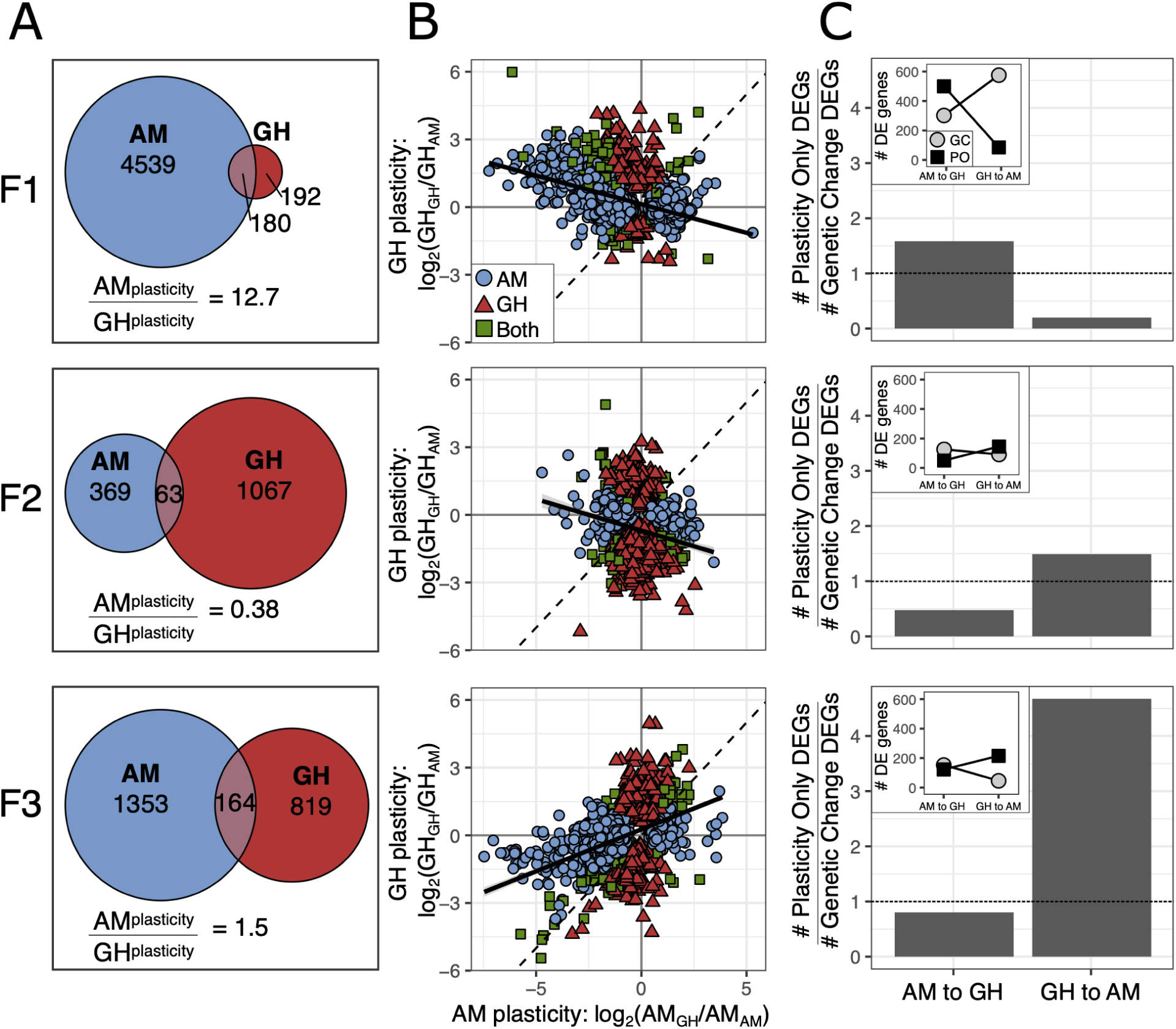
Plasticity in gene expression in response to reciprocal transplant across three generations in transplant conditions. (A) Gene expression plasticity for AM and GH transplanted lines as defined by differentially expressed genes within a line following transplant (GH_GH_ vs. GH_AM_; AM_AM_ vs. AM_GH_). (B) Comparison of plastic changes between AM and GH lines. Color and shape indicate line. If plastic responses are equal between the two lines, the slope of the relationship would be 1 (dashed black line). The observed slope is shown as the solid black line. (C) Plastic versus genetic changes in gene expression. Bar plots show the relative number of plastic versus genetic changes in expression between AM_AM_ and GH_GH_ where the dashed line indicates equal numbers. AM to GH indicates AM lines moving to GH conditions, GH to AM is the opposite. Inset plots show the counts of the genes that went into the barplot where PO = plasticity only and GC = genetic change. All proportions of PO to GC significant at *P* < 0.001, G test of independence.

To understand the relative expression patterns of the plastic genes, we compared gene expression plasticity in GH versus AM lines in response to each environment (measured as log_2_ fold change: AM_GH_ vs. AM_AM_ and GH_GH_ vs. GH_AM_; Fig. 3B). When comparing expression patterns in response to environment, the expectation for equal transcriptional plasticity and equal response to environment for GH and AM lines would be a slope of one among genes differentially expressed. If there was a reduction in plasticity, perhaps due to genetic assimilation, the evolved line (GH) would have an expression response with a slope less than one. After one generation in transplant conditions, there was a slight negative relationship, slope of -0.25 (Fig. 3B). A negative relationship between expression responses could be interpreted as non-adaptive gene expression ^19^ or as a consistent response to transplant between the lines, i.e., a shared stress response to transplant and an opposite response to environmental condition. In the latter case, equal gene expression and plasticity in response to transplant between the lines would yield a slope of -1. In the second generation of transplant, the negative relationship persisted (slope = -0.29), but by F3 there was a positive relationship where each transplanted line converged on the gene expression profile of its new environment (slope = 0.37). However, the slope was less than 1, indicating that AM lines had a greater change in expression by F3 than GH lines and were better able to match their new environment. While expression changes at F1 were likely due to plasticity, differential gene expression at F2 and F3 could be due to plasticity or evolution post-transplant. For example, transgenerational plasticity may cause the gene expression at F1 to be influenced by the environment of the previous generation ^42, 64, 65^. Additionally, genetic adaptation across the generations could drive a shift in gene expression patterns following transplant ^41^. Each of these explanations would support the pattern of successive changes across each generation and we disentangle their relative impacts below. Regardless of the mechanism, the loss of plasticity with adaptation to the higher temperature of the GH environment matches predictions from natural populations of *A. tonsa*. Populations sampled along a latitudinal gradient show a trade-off between thermal tolerance and degree of developmental plasticity ^66^. The same pattern is observed in *A. tonsa* populations throughout the season ^67^.

Gene expression patterns after long-term experimental evolution and reciprocal transplantation also allow us to test the hypothesis that transcriptional plasticity can be retained and serve as a mechanism to facilitate re-adaptation to ancestral conditions in the context of adaptation to global change conditions and extreme events. Using the approach described in Ho et al. ^68^, first we identified the genes differentially expressed between AM_AM_ and GH_GH_, changes likely to be adaptive in each environment. These genes were further characterized as requiring “plasticity only” or requiring “genetic change” for both forward adaptation, AM to GH conditions, and reverse adaptation, GH to AM conditions, by comparing the relative expression levels in home versus transplant environments. Specifically, for the forward direction, genes were considered plasticity only (PO) if they were differentially expressed between AM_AM_ vs. AM_GH_ and not differentially expressed between AM_GH_ vs. GH_GH_. Genes were considered to require genetic change (GC) if they were differentially expressed between AM_GH_ vs GH_GH_. PO and GC genes were mutually exclusive but did not include all of the genes differentially expressed between AM_AM_ and GH_GH_ (F1: 87% of total; F2: 66%; F3: 80%). Last, we similarly categorized the differentially expressed genes as PO or GC for the reverse adaptation from greenhouse to ancestral ambient conditions: PO: differentially expressed between GH_GH_ vs. GH_AM_ and not differentially expressed between GH_AM_ and AM_AM_; GC: differentially expressed between GH_AM_ vs. AM_AM_.

After 20 generations of experimental evolution and one generation of reciprocal transplantation, forward adaptation had more plasticity-only genes (Fig. 3C; 537 genes) than genetic-change-needed genes (Fig. 3C; 339 genes). In contrast, reverse adaptation, i.e., returning to ambient conditions after 20 generations of greenhouse adaptation, showed the opposite pattern with substantially more genetic change needed (Fig. 3C; 613 genes) than plasticity genes (Fig. 3C; 122 genes). These results suggest that forward adaptation to greenhouse conditions was likely initially possible using plasticity (in this case, the plasticity was maintained in the ambient control lines), and that reverse adaptation, after 20 generations in constant conditions, had little plasticity remaining and required genetic change. These results are in contrast to what Ho et al. ^68^ found in high-altitude adapted chickens, natural and introduced guppy populations, and experimental evolution in *E. coli* where genetic change genes were more common when moving to the new environment and plasticity-only genes were more common when going back to the original environment. After two and three generations of transplant, however, our results match Ho et al.’s patterns: reverse adaptation, GH in AM, had more plasticity-only genes than genetic change needed genes, while forward adaptation, AM in GH, had more genetic change needed than plasticity only genes (Fig. 3C). It is important to note that F2 and F3 animals had been in the transplant environment for 2 and 3 generations, respectively; gene expression responses could be due to transgenerational plasticity or rapid evolution post-transplant. Hence, we show that phenotypic plasticity does not always ease adaptation back to ancestral environments. Instead, it can facilitate adaptation to a novel environment and be rapidly lost, in accord with theoretical work ^17^, and recover via transgenerational or genetic mechanisms, explored next.

To explore the relationship between gene expression and allele frequency changes across transplant generations, we used Discriminant Analysis of Principal Components (DAPC). Discriminant function space was generated from AM_AM_ and GH_GH_, representing the adaptive expression differences after 20 generations in each environment (Fig. 4, blue and red shaded zones). Transplanted lines were fit to this discriminant function space to determine if and how expression changed to match the adaptive expression profile where the adaptive plasticity is summarized by the length of each arrow in Fig. 4A. Matching the results above, we found lower overall plasticity in GH transplanted copepods relative to AM transplanted copepods at F1 (Markov chain Monte Carlo (MCMC) generalized linear mixed model; *P_MCMC_* = 0.03; Fig. 4A). However, by F2 and F3, AM and GH lines demonstrated similar shifts where lines converge on the adaptive expression profile by the F3 generation (*P_MCMC_* > 0.05; Fig. 4A). Alternative methods based on DeSeq2 differential expression recapitulate these sequential adaptive expression shifts (Fig. S5). This increase in adaptive gene expression across generations provides additional evidence that copepods, even after 20 generations of experimental evolution in ambient or greenhouse conditions and despite differences in initial levels of plasticity, were able to match adaptive gene expression profiles increasingly with each successive generation.

**Figure 4.**
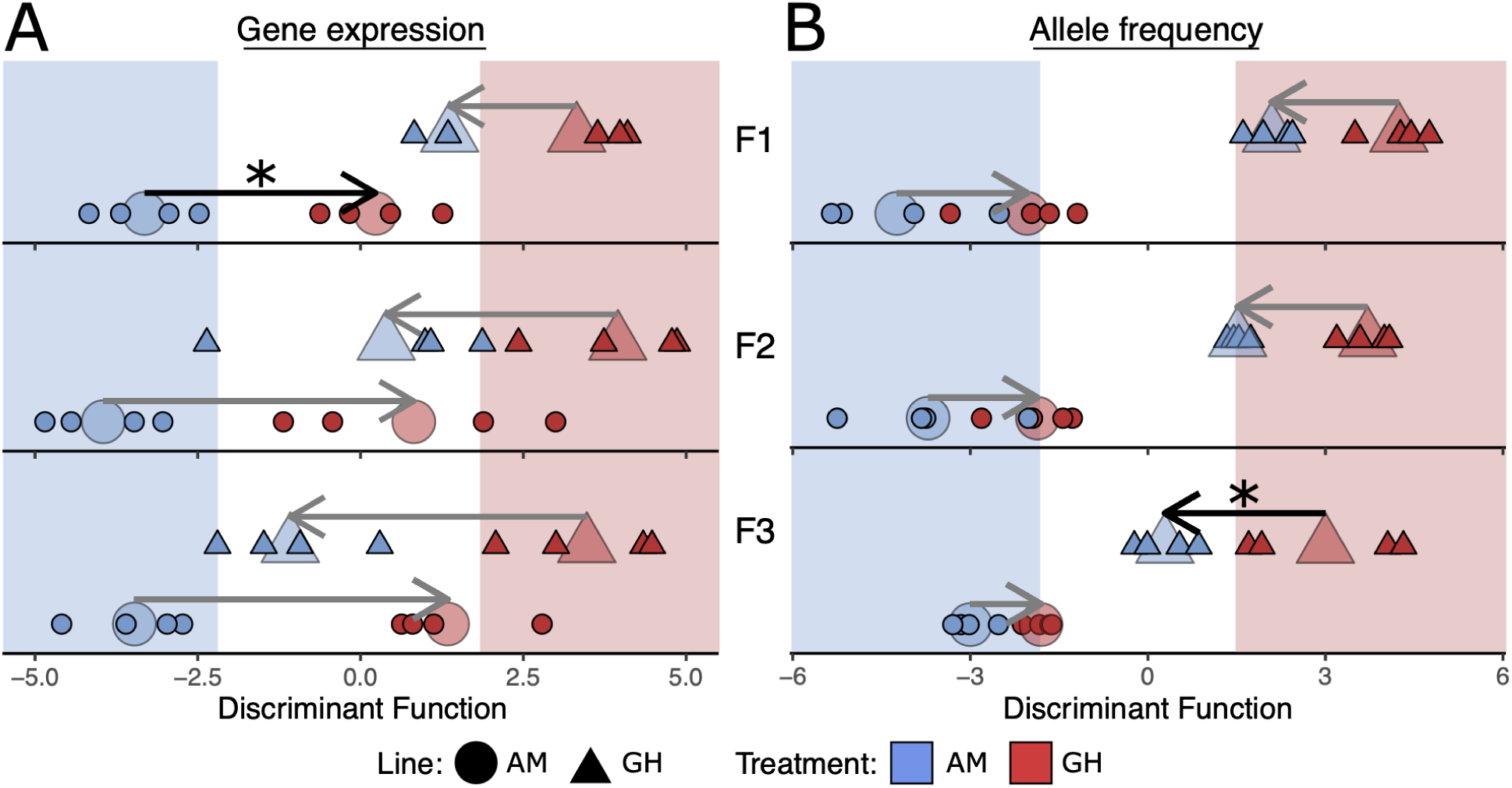
Genome-wide variation in (A) gene expression and (B) allele frequencies across transplant generations. The x-axis shows the discriminant function space that maximizes differences between lines in their home environments and the background shading represents the “home” discriminant function space for each line. Shape and color of points indicate selection line and treatment condition, respectively. Small points are individual replicates while the mean change for each group is represented by the large transparent points; arrows connect home to transplant means. Significantly different shifts between lines are represented by black arrows and asterisks. Gene expression profiles of transplanted lines converge on the expression of the non-transplanted counterpart in their new environment to a greater extent for AM as compared to GH lines at F1 (P_MCMC_ = 0.03) but lines show similar movement at F2 and F3 (P_MCMC_ > 0.05). Conversely, allele frequencies change to a greater degree in the GH line than the AM line by generation F3 (P_MCMC_ = 0.02).

Using a similar DAPC approach with allele frequencies, we tracked shifts in allele frequencies towards adaptive alleles following transplant for each successive generation. At generations F1 and F2, we found allele frequency changes were similar between AM and GH lines (*P_MCMC_* > 0.05; Fig. 4B). However, by F3 transplanted GH animals showed significantly greater adaptive evolution than AM (*P_MCMC_* = 0.02). These findings were again mirrored by an independent analysis based on Cochran–Mantel–Haenszel tests (Fig. S5). Together with the gene expression analysis, these results indicate that AM_GH_ animals were able to match the greenhouse adaptive transcriptional profiles through physiological and/or transgenerational plasticity and minor changes in allele frequencies. Conversely, the GH_AM_ animals had lost the plasticity necessary to transcriptionally respond and matched the adaptive transcriptional profile through evolution, i.e., selection drove changes in allele frequencies across the three generations of transplant.

Rapid adaptation can carry costs at the genetic and phenotypic levels. For example, a selective bottleneck can drive a loss of genetic diversity during long-term selection or short-term transplant. To test for this possibility, we quantified nucleotide diversity (π) for each treatment following 20 generations of selection and after transplant. We found no significant loss of genetic diversity after long-term adaptation to greenhouse conditions (GH_GH_, Fig. 5A, *P_Tukey_* > 0.05). Similarly, ambient lines, both transplanted and not, maintained high levels of genetic diversity (*P_Tukey_* > 0.05). However, when transplanted back to ambient conditions, GH_AM_ showed a significant drop in diversity by F3 (*P_Tukey_* < 0.05; Fig. 5A), indicating a global loss of genetic diversity. This suggests that the successive shifts in allele frequency after transplant of GH to AM (GH_AM_; Fig. 4B) resulted in a loss of genetic diversity (Fig. 5A). Taken together, these results indicate that matching gene expression to the environment after transplant (GH to AM) was driven by adaptation across multiple generations rather than plasticity; this loss of plasticity is evidence of genetic assimilation. To test if this loss of genetic diversity in transplant from GH to AM was concentrated in specific regions of the genome, we compared changes in nucleotide diversity in regions identified as adaptively diverged between greenhouse and ambient lines (i.e., regions containing 17,720 consistently divergent loci) versus all other loci (“non-adaptive”). For GH_AM_ lines, the loss of nucleotide diversity was concentrated in genomic regions containing adaptive loci (Fig. 5B; Wilcoxon test *P* = 0.007), while there was no loss of diversity in AM_GH_ or GH_GH_ lines as control contrasts (Fig. 5B; *P* = 0.16; *P =* 0.07). In addition, the loss of nucleotide diversity for GH_AM_ lines was concentrated in genes with functions related to sequestration of actin monomers, cytokinesis, and response to stress (Fig. 5C; Table S3), similar to those underlying adaptive genetic divergence between AM_AM_ and GH_GH_ lines (Fig. 2; Table S1). AM_GH_ and GH_GH_ showed no functional enrichment for regions losing genetic diversity. The lack of nucleotide diversity loss and the absence of functional enrichment indicates that selection did not act on AM_GH_ or GH_GH_ transplanted lines, which is consistent with the sustained ability for a plastic response in AM lines but a loss of plasticity in GH lines.

**Figure 5.**
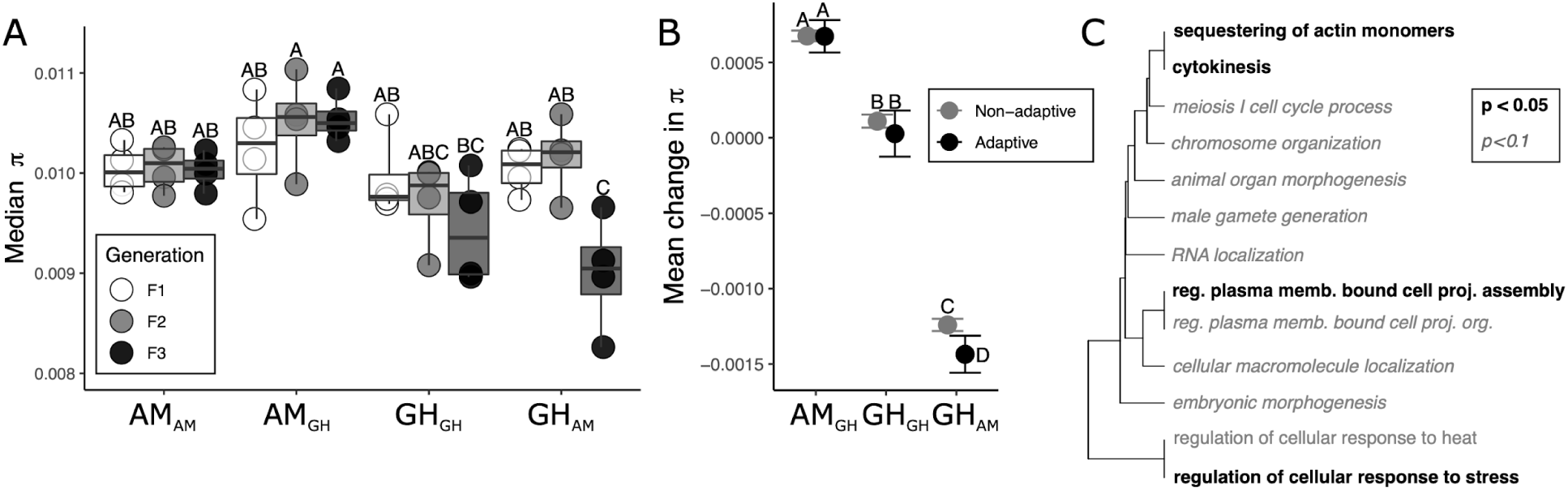
Changes in genetic diversity. (A) Median nucleotide diversity (π) for all treatments and generations following transplant. Boxes represent the upper and lower 25% quantiles and median while whiskers are min and max. (B) Mean change in π from after transplant for adaptive and non-adaptive SNPs with 95% confidence intervals. Changes relative to AM_AM_ F1. Letters above each point (for A and B) show significance; points sharing letters are not significantly different. (C) Gene ontology enrichment for the loss of genetic diversity of GH_AM_ at transplant F3; AM_GH_ showed no enrichment.

To identify life-history traits that could be under selection and reveal potential costs of adaptation, we measured egg production rate across three days of adulthood, survivorship from nauplii to reproductive adult, and combined these metrics with development rate, time to adulthood, and sex ratio to estimate net reproductive, lambda ^69^ in the first generation after transplant (F1). We found that fecundity was equally high for both lines in their home environments but declined for GH lines after transplant to AM conditions (48% reduction in egg production relative to GH_GH_ ; *P* < 0.02; Fig. 6A). Conversely, transplanted AM_GH_ were able to maintain similarly high egg production as the non-transplanted lines (Fig. 6A). In contrast to egg production, survivorship from *nauplii* to reproductive maturity was 60% lower for greenhouse compared to ambient lineages (*P* < 0.001; Fig. 6B). Similar to egg production rates, lambda, reflected stable population sizes for AM lines in both conditions and GH_GH_, but declined with GH_AM_ transplant (*P* = 0.01, Fig. 6C); these estimates were congruent with observed culture densities. The estimated stable growth rate of GH_GH_ lines despite decreased survival relative to AM lines (Fig. 6B) was due to compensation in other life-history traits including higher egg production, faster development, and earlier reproductive timing in GH conditions ^70^, as has been observed in many other taxa including *Drosophila* ^71^. These results suggest that egg production rate, faster development and earlier time to adulthood may have been life-history traits diverging between ambient and greenhouse conditions, yielding stable population sizes to compensate for relatively poor survival. That poor survival was not observed for AM animals in GH conditions suggests GH lines lost plasticity and that there was not a cost to maintaining this plasticity in the constant AM conditions.

**Figure 6.**
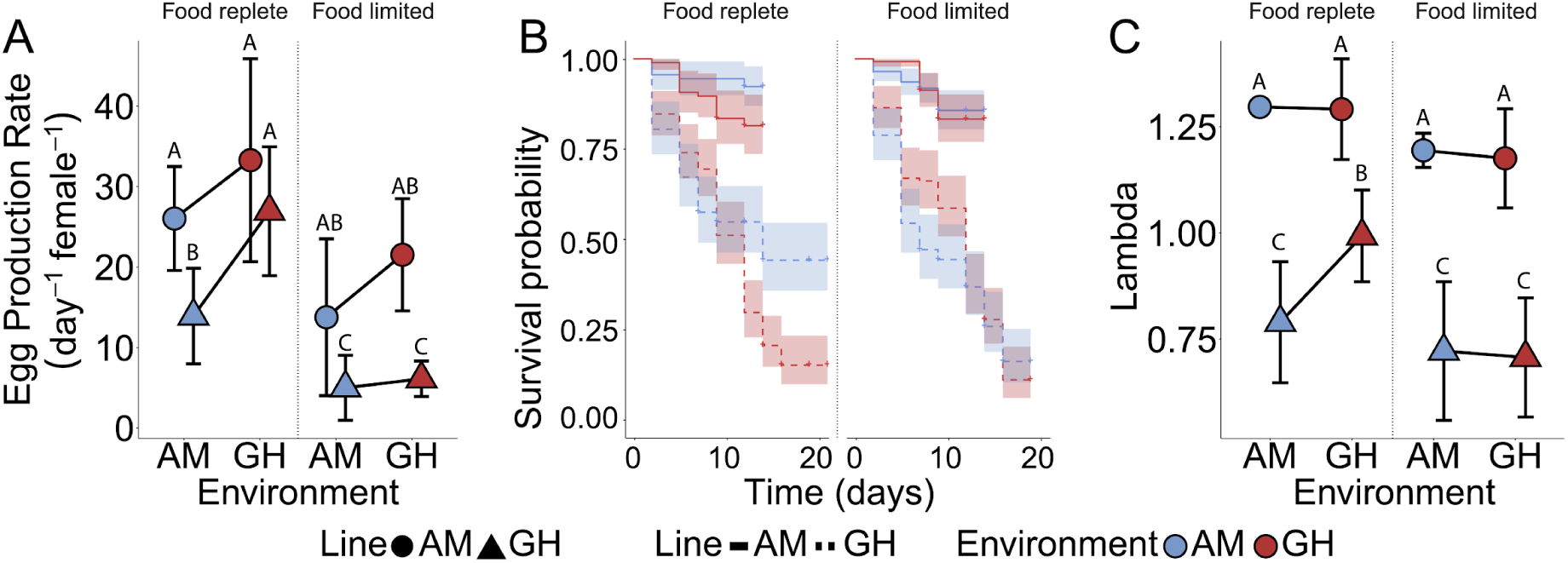
Life-history traits after 20 generations of selection in ambient (AM) and greenhouse (GH) conditions and one generation of reciprocal transplant. For all panels, shape indicates selection line, color indicates environmental condition, and error bars represent 95% confidence intervals. A) Egg production rate under ad-libitum and food-limited conditions. B) Survivorship under ad-libitum and food-limited conditions. C) Lambda (net reproductive rate) calculated from combined fecundity, development rate, sex ratio, and survivorship data where a values greater than 1 indicate positive growth. Capital letters in A and C represent statistical similar groups.

The lower survivorship and lambda of the GH lines relative to AM_GH_ transplants suggests that evolution in GH conditions occurred with a trade-off among life history traits, e.g., survival and age at reproduction ^72, 73^ or a trade-off at the physiological level. In experimental evolution studies, lower performance of stress-adapted lines in ancestral environments is common ^74–76^, particularly for complex organisms and when stressful conditions are constant ^77–79^. The reduced ability of GH-adapted lines to maintain high fitness across environments could be due to antagonistic pleiotropy, trade-offs at the molecular, cellular or life-history trait levels, via selection for a locus associated with multiple traits which are adaptive in one environment and mal-adaptive when the environment changes ^80^.

To test the hypothesis that adaptation to a stressful environment could reduce fitness when conditions change to a novel stressor, we reared all evolved and transplanted lines in low food conditions, a stressor that is predicted to increase in frequency as climate changes ^81^, and measured the same life-history traits at F1. In support of this hypothesis, we found that GH animals performed poorly under low food availability regardless of environment (Fig. 6). Under low food, relative to AM lines, GH lines experienced a 67% reduction in egg production rates (*P* < 0.001; Fig. 6A), 70% reduction in survival (*P* < 0.001; Fig. 6B), and a 40% reduction in fitness (*P* < 0.001; Fig. 6C). These results suggest that selection in greenhouse conditions may result in populations which are less resilient to further environmental changes, including expected changes such as a return to ambient conditions or low food availability ^82^. This reduced flexibility may be driven by increased energetic costs of life in greenhouse conditions leading to increased stress under low food conditions ^83, 84^ or antagonistic pleiotropy and the costs of adaptation for a complex organism to adapt to a complex stressor ^79, 85^, both high pCO_2_ and high temperature. In the case of increased metabolic costs in GH conditions, we would not expect genetic divergence between the lines and would see limited differences in plastic transcriptional responses to transplant; the loss of robustness in GH animals would be due to lower habitat quality and not due to directional selection. Instead, we observe a signal of decreased plasticity driven by selection where there was consistent genomic divergence between GH and AM lines coupled with a high degree of transcriptional plasticity in AM animals that is not present in GH animals. This result supports previous findings that long-term evolution in a constant, stressful selective environment can result in specialization, i.e., a loss of plasticity, for example due to antagonistic pleiotropy ^74, 78^ or due to high costs of maintaining a generally plastic phenotype ^86^. Alternatively, this rapid loss of plasticity could be driven by neutral mechanisms following the relaxation of selection for plastic phenotypes in a static environment, such as drift or the accumulation of mutations, though the relatively short time frame makes this less likely than selective processes ^87^. Under either mechanism, in the context of global change, such specialization could rapidly occur for populations of a short-lived species experiencing an extreme event.

## Conclusion

Combining long-term experimental evolution of a marine metazoan with three generations of reciprocal transplant, we reveal important interactions between plasticity and genetic adaptation in response to global change conditions. In the short-term, three generations, plasticity buffered populations from environmental change allowing AM lines to tolerate greenhouse conditions through plasticity. Our data suggest that over the longer-term, 20 generations, plasticity to acclimate was lost as adaptation to greenhouse conditions proceeded, indicating costs to maintain plasticity ^86^. Our results advance the understanding of the relationship between plasticity and adaptive evolution by providing evidence that plasticity did not impede adaptation but, over time, adaptation eroded plasticity. Importantly, by including multiple generations of transplantation following 20 generations of selection, we show that, even with a loss of plasticity, animals can re-evolve back to their ancestral conditions. However, this re-adaptation carries the cost of losing adaptive genetic variation.

This interplay between plasticity and adaptation has important implications for our understanding of mechanisms of species persistence to global change conditions. We demonstrate that copepods have the capacity to adapt over 20 generations, approximately one year, using the standing genetic variation that exists in natural populations. However, reciprocal transplant and food challenges revealed significant costs of this adaptation; populations lost physiological plasticity, the ability to tolerate food limitation, and ultimately adaptive genetic variation for greenhouse conditions. Given the increasing frequency of extreme environmental conditions, rapid adaptation of *A. tonsa* to a temporary extreme may result in a population that is maladapted when conditions return to the previous state. Thus, while plasticity may enable persistence initially, adaptation can drive a loss of plasticity that can lead to reduced resiliency as environmental conditions continue to fluctuate. These results demonstrate the utility of experimental evolution in understanding complex adaptation in natural systems and to reveal the mechanisms, time-scales of responses, consequences, and reversibility of this process. As we begin to incorporate adaptive potential and plasticity into species persistence models ^88–91^, these results caution that, even for species predicted to be resilient to rapid global change, there may be unforeseen costs to adaptive evolution.

## Methods

### Experimental set-up

Copepods (n=1,000) were collected in June of 2016 from Esker Point Beach in Groton, CT, USA (decimal degrees: 41.320725, -72.001643) and raised for at least three generations prior to the start of selection to limit maternal effects ^65^. Parental individuals were acclimated to one of two experimental conditions: 1) AM – Ambient temperature (18 °C), Ambient CO_2_ (pH ∼ 8.2; pCO_2_ ∼ 400 μatm) or 2) GH – High temperature (22 °C), high CO_2_ (pH ∼7.5; pCO_2_ ∼ 2000 μatm; see table S7 for measured values). Cultures were fed every 48-72 hours at food-replete concentrations of carbon (>800 μg C/L) split equally between three species of prey phytoplankton. Prey phytoplankton included *Tetraselmis* spp., *Rhodomonas* spp., and *Thalassiosira weissflogii*, which is a common diet combination used for rearing copepods ^92^. About 400 females and 200 males were used to seed each treatment, yielding an estimated 15,000 eggs and newly hatched *nauplii* for the initial F0 generation. Four replicate cultures per treatment were maintained in individual 3L culture containers where each temperature was housed in one of two identical incubators: one incubator for GH conditions, one for AM conditions. Elevated CO_2_ levels were achieved with two-gas proportioners (Cole-Parmer, Vernon Hills, IL, USA) combining air with 100% bone dry CO_2_ that was delivered continuously to the bottom of each replicate GH culture. Actual measurements of experimental CO_2_ were determined based on measurements of salinity, temperature, pH, and total alkalinity using CO_2_Sys. Total alkalinity was measured before and after the course of the experiment. Seawater was filtered to 30 μm and measured immediately using an endpoint titration as described previously ^93^. Target pH values were monitored daily using a handheld pH probe (Orion Ross Ultra pH/ATC Triode with Orion Star A121 pH Portable Meter (Thermo FisherScientific®, Waltham, MA, USA) which was calibrated monthly using commercially available pH standards in a three-point calibration (pH 4.01, 7, 10.01; Thermo FisherScientific®; Waltham, MA, USA). Continuous bubbling maintained higher than necessary dissolved oxygen levels (>8 mg/L). To assess functional life-history traits, smaller volume experiments were housed in the same temperature-controlled incubators in custom plexiglass enclosures with the atmosphere continuously flooded with CO_2_ infused air at the appropriate concentration, which allows for passive diffusion of CO_2_ into the experiments. Copepods were raised in the respective two experimental treatments for 20 generations before the reciprocal transplant. To create non-overlapping generations, at each generation, adults were removed after laying eggs to allow for tracking of each generation. Adults were left to lay eggs for 1 week after we observed the first evidence of a new generation (i.e. new *nauplii*). This allowed nearly all surviving adults to procreate while minimizing the chance that remaining adults would cannibalize or mate with their offspring.

### Reciprocal transplant

At the 21st generation, we performed a reciprocal transplant between copepods from AM and GH. Each lineage x environment interaction yielded four unique treatments. Each of the four replicates from each treatment was split to yield four additional replicates for each of two new transplant treatments: AM_GH_ and GH_AM_ (as well as sham transfers: AM_AM_ and GH_GH_). Transplant treatments were housed in incubators that corresponded to the new environments along with the original treatments: AM_GH_ and GH_GH_ were housed in the GH incubator while GH_AM_ and AM_AM_ cultures were housed in the AM incubator. Copepods were raised for three subsequent generations and were fed every 48-72 hours as described above. This design led to 48 total samples for genomics: 2 treatments × 2 transplant/non-transplant × 4 replicates × 3 generations.

### Genomics

RNA from pools of 20 individuals was extracted using TRIzol reagent (Invitrogen, Carlsbad, CA, USA) and purified with Qiagen RNeasy spin columns (Qiagen, Germantown, MD, USA). RNAseq libraries were prepared by Novogene (Sacramento, CA, USA) and sequenced with 150 bp paired end reads on an Illumina NovaSeq 600, generating 1.29 billion reads. Raw reads were trimmed for quality and adapter contamination using Trimmomatic V0.36^94^ where leading and trailing quality was set at 2, sliding window length was 4 with a quality of 2, and minimum length was 31.

To quantify genetic variation, SuperTranscripts were generated from the reference transcriptome ^95^ using the Trinity_gene_splice_modeler.py command in the Trinity pipeline ^96^. SuperTranscripts act as a reference by collapsing isoforms into a single transcript, which enables variant detection without a reference genome ^97^. Trimmed reads were aligned to the SuperTranscripts using bwa mem ^98^ and duplicate reads were marked with Samblaster ^99^.

Variants were called using VarScan v2.3 ^100^ (2 alternate reads to call a variant, p-value of 0.1, minimum coverage of 30x, minimum variant allele frequency of 0.01). Following this, variants were stringently filtered to include no missing data, only sites where at least 4 samples were called as variable and each with a minor allele frequency of at least 0.025 (minimum of 1 heterozygous individual), any sites with depth greater than 3x the median (median=132x), and sites with a minimum coverage of 50x per sample (minimum 2.5 reads/diploid individual; average 174× coverage, average 8.7 reads/diploid individual after filtering; Fig. S6). This reduced called variants from an initial 5,547,802 to 322,595.

Gene expression was quantified at the gene level as recommended ^101^ using Salmon v0.10.2 ^102^ and transcript abundances were converted to gene-level counts using the tximport package in R ^103^. The reference transcriptome was indexed using quasi mapping with a k-mer length of 31 and transcripts were quantified while correcting for sequence specific bias and GC bias. We were interested in assessing expression patterns between treatments with each generation and how these patterns changed through time and, therefore, analyzed each generation separately. Genes with low expression (fewer than 10 counts in more than 90% of the samples) were removed from the dataset. This left 23 324, 24 882, and 24 132 genes for F1, F2, and F3, respectively. Counts were normalized and log transformed using the rlog command in DESeq2 ^104^.

### Gene expression analysis

DESeq2 was used to identify differentially expressed genes. For each generation, we required at least 10 counts in 90% of samples. Differentially expressed genes were identified using the model ∼Line + Treatment + Line:Treatment and any contrasts were considered significant when adjusted p-values were < 0.05. We identified the plastic genes for each line as the genes changing expression between environments within a line (GH_GH_ vs. GH_AM_; AM_AM_ vs. AM_GH_). We compare the plasticity between AM and GH lines by plotting the log_2_ fold change between GH_GH_/GH_AM_ and AM_GH_/AM_AM_, similar to approaches taken in other plasticity research ^18, 21^. To assess the mechanisms driving gene expression divergence between lines in their home environments after 20 generations, we partitioned gene expression changes into Plastic Only and Genetic Change genes ^68^. For genes that were significantly different between GH_GH_ and AM_AM_ (F1: 1129 genes; F2: 527 genes; F3: 465 genes), we assessed the mechanisms enabling the transition from AM to GH (forward evolution) and GH to AM (reverse). For AM to GH, when a gene was differentially expressed between the transplanted and home line (AM_GH_ vs. GH_GH_), the gene expression shift required genetic change. Alternatively, if the gene was significantly plastic (significant AM_AM_ vs. AM_GH_) with no genetic change, this was a plasticity only gene. These two categories were mutually exclusive but did not include all genes differentially expressed between AM_AM_ and GH_GH_ due to the cases where genes were not differentially expressed between AM_AM_ vs. AM_GH_ or AM_GH_ vs. GH_GH_, even though gene expression was different between AM_AM_ vs GH_GH_. We conduct the opposite analysis for GH to AM. With this analysis we can partition the changes in expression after 20 generations of selection into those that were achieved through plastic versus evolved mechanisms.

### Discriminant analysis of principal components

Discriminant analysis of principal components (DAPC) was used to quantify the degree to which gene expression and allele frequencies converged on the adaptive state within each environment. Here, we assume that each line has reached the adaptive optimum after 20 generations. Using the filtered, normalized, and transformed expression data from DeSeq2, shifts in gene expression in transplanted lines across generations were quantified using DAPC in the adegenet package in R ^105^. Discriminant functions for each generation were first generated using the non-transplanted lines to identify genes consistently differentially expressed. Four principal components that represented 81, 78, and 80% of the variation at F1, F2, and F3 were retained and two clusters that represented the two non-transplanted lines were used. Transplanted lines were fit to this discriminant function space and used MCMCglmm models in R ^106^ to model the effect of line origin and transplant on movement in discriminant function space. 2500 posterior estimates were generated and the difference in the transplant effect for each line was quantified by calculating the absolute difference between these estimates. This can be viewed as the difference between the lengths of the lines representing average shifts in Fig. 4. These differences were used to generate a 95% credible interval and the proportion of positive or negative values were considered a p-value for the difference in magnitude of the effect of transplant on each line in discriminant function space.

DAPC was also used to identify divergence in allele frequency between the AM and GH lines. The same approach as for gene expression (above) was taken where we generated the DAPC with lines in their home environment and then fit transplanted lines to discriminant function space. Here, we retained three principal components that represented 56, 59, and 63% of the total variation at each generation and again used two clusters.

### Adaptive allele frequency and gene expression divergence

To identify loci consistently shifting in response to selection, Cochran–Mantel–Haenszel (CMH) tests were used to identify specific SNPs that were consistently diverged between the AM and GH lines and represent the most likely targets of adaptation. This approach looks for consistent changes in allele frequencies across all replicates and is a common and powerful technique in experimental evolution ^24, 107^. We calculated CMH p-values for AM_AM_ versus GH_GH_ lines (n = 4, each) for each of the three sampled generations. Significance thresholds (*P* < 5.17e-08) were defined with Bonferroni corrections using the total number of SNPs (322,595) multiplied by three, representing the total number of tests conducted. The SNPs that were identified in all three generations were considered adaptive (Fig. S7). This approach assumes that adaptive differences between AM_AM_ and GH_GH_ lines after 20 generations are much larger than any additional response to selection during the three generations of transplant. As such, the signals of adaptation in the home environment at F1 should be consistent in F2 and F3 and can be used to identify loci under selection.

DESeq2 was used to identify the specific genes that were adaptively differentially expressed between non-transplanted AM and GH lines at each generation. Using the same geneset and model from DeSeq2, above, we identified differentially expressed genes between AM and GH in their home environment with an adjusted significance threshold of p-values < 0.05.

### Quantifying genetic diversity

We estimate genetic diversity (π) for each replicate using Popoolation ^108^ with 100 bp non-overlapping sliding windows. To quantify if π was differentially lost in any treatment, median π values were compared using an Anova with a Tukey post-hoc test. We next tested if regions containing adaptively divergent loci between non-transplanted GH and AM lines lost π at a different rate than regions containing only neutral variants. The change in π was calculated for each replicate and the change in π for GH_AM_ and AM_GH_ were compared using Wilcoxon Rank Sum tests with bonferroni corrections. Gene ontology enrichment was performed with GO Mann-Whitney U ^109^, which requires no significance threshold, but takes the change in π for each group and asks if any functional category falls towards the tails of this distribution (in our case, one-sided to identify disproportionately low values). We use this same approach for gene expression and allele frequencies divergence between lines in their home environments. For allele frequencies, the minimum p-value was chosen for each gene.

Finally, we tested for overlap between the allelic and gene expression results to determine if the genes in each were significantly correlated. We correlated the relationship between the -log10 of the CMH p-values for the allele frequencies and the differential gene expression p-values between GH_GH_ and AM_AM_. This analysis showed the two sets were distinct and the variation explained by each was less than 1% at each generation, indicating that there was minimal bias in estimating allele frequencies due to differentially expressed genes.

### Life-history traits

Day-specific survivorship was measured every 48-72 hours, with food media replaced on monitoring days. Food media was provided at the appropriate food concentration (>800 μg C/L for food-replete, 250 μg C/L for food-limited, and 0 μg C/L for starved) divided in equal carbon proportions between the three afore-mentioned prey species in 0.2 μm filtered seawater collected from Long Island Sound. Food media was acclimated to the appropriate temperature and CO_2_ concentration prior to replacement. Survivorship was assessed among twelve 250-mL beakers per treatment (3 food concentrations × 4 replicates = 12 beakers per treatment) containing 25 individual N1 *nauplii* in the same plexiglass enclosures as described above and monitored until sexual maturity (adulthood). Life-history results are only presented for food replete and food limited conditions because no starved individuals survived. Log rank analysis of survivorship was assessed using the survival ^110, 111^ and survminer ^112^ packages in R.

Egg production rate (EPR) and hatching success (HS) were assessed with 36 individual mate pairs of newly matured adults per treatment (3 pairs per food concentration × 3 food concentrations x 4 replicates = 36 pairs per treatment). Adults were incubated in 25 mL petri dishes (FisherScientific, Waltham, MA, USA) over three days in the same temperature-controlled incubators and plexiglass enclosures described above. After the initial three-day incubation, adults were assessed for survival and removed to avoid cannibalism of eggs. Eggs were allowed to hatch over a subsequent three-day incubation. Food media was prepared as described for survivorship and replaced daily during egg laying to ensure accurate food concentrations were near saturation and not reduced due to daily grazing ^113^. Lids of petri dishes were left off-center to allow for full contact with the atmosphere and diffusion of CO_2_. Plates with dead males were still evaluated for EPR, but not HS. Plates with dead females were not evaluated because we cannot estimate egg production or hatching when no females were alive; mortality was estimated separately and we do not want to confound these two measures. The maximum number of plates excluded was six in the GH_AM_ treatment at food limited conditions. No other treatment experienced more than three plates excluded from the assay. After the hatching period, plates were preserved with non-acid Lugol’s solution and eggs counted and nauplii counted. Per capita EPR was calculated as (E_u_+E_h_)/t where E_u_ represents unhatched eggs, E_h_ represents hatched eggs (*nauplii*), and t represents egg laying time. Hatching success was calculated as E_h_/(E_u_+E_h_).

The population net reproductive rate, λ, was calculated as the dominant eigenvalue of an assembled projected age-structured Leslie Matrix constructed from survivorship and fecundity data ^69^. Briefly, day-specific probabilities of survivorship are calculated from day-specific survivorship as 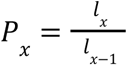 represents the number of individuals on day *x* and *l* _*x*−1_ *_x_* represents the number of individuals on day *x-1*. Probabilities of survivorship on day 1 are assumed to be 100%.. Per capita EPR and HS are calculated as described above, with fecundity rates equaling the product of EPR and HS. Because only females produce offspring, total fecundity rates must be scaled to the sex ratio (ratio of females:males) observed in survivorship experiments. To account for differences in individual development time for each treatment, fecundity rates are assigned to all days after the first matured adult is observed. We assume that survivorship in each beaker is equally as likely to experience the fecundity values observed in EPR experiments. Therefore, each mate-pair fecundity rate was paired with each survivorship beaker to construct a matrix. This yields a maximum of 48 matrices per treatment per food concentration (4 beakers × 12 mate pairs). Errors of lambda were calculated as the 95% confidence interval using the formula: 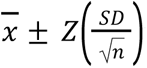 where 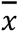 represents the mean, *Z* represents the 95% z-score (1.960), SD represents the standard deviation of the mean, and n represents the number of observations.

We constructed linear mixed models with line, environment, and food concentration as fixed effects with all interactions, and culture replicate as a random effect. The results for net reproductive rate fall within a normal distribution with an inflation of zero values. Thus, we constructed zero-inflated generalized linear mixed models to account for the additional zeroes. Two post-hoc tests were conducted. First, to quantify differences in plasticity between the lines, we evaluated genotype × environment interactions on life-history traits separately for each food concentration. Second, post-hoc t-test comparisons were used to conduct pairwise comparisons of the interactions for each life history trait.

## Acknowledgments

We thank Lydia Norton and Gihong Park for culture maintenance and experimental help. RSB appreciates the sponsorship from Maile Neel as a Visiting Research Scientist in the Department of Plant Science & Landscape Architecture at the University of Maryland College Park.

## Author contributions

Conceptualization and experimental design: all authors; Experimental execution: JAD; Data collection and analysis: RSB and JAD; Writing-original draft: RSB and MHP; Writing-review and editing: all authors

## Data and materials availability

Raw sequence data is available at NCBI BioProject PRJNA555881. Life history data are available as a supplemental file and allele frequency and gene expression table is available on Figshare (https://doi.org/10.6084/m9.figshare.10301690). Correspondence and requests for materials should be addressed to RSB or MHP. Code to reproduce all analyses can be found at: https://github.com/PespeniLab/tonsa_reciprocal_transplant

## Extended data

Figures S1-S8

Tables S1-S7

